# Changing planetary rotation rescues the biological clock mutant *lhy cca1* of *Arabidopsis thaliana*

**DOI:** 10.1101/034629

**Authors:** Andrew J. Millar, Jamie T. Carrington, Wei Ven Tee, Sarah K. Hodge

## Abstract

**Background**: Pervasive, 24-hour rhythms from the biological clock affect diverse biological processes in metabolism and behaviour, including the human cell division cycle and sleep-wake cycle, nightly transpiration and energy balance in plants, and seasonal breeding in both plants and animals. The clock mechanism in the laboratory model plant species *Arabidopsis thaliana* is complex, in part due to the multiple interlocking, negative feedback loops that link the clock genes. Clock gene mutants are powerful tools to manipulate and understand the clock mechanism and its effects on physiology. The *LATE ELONGATED HYPOCOTYL* and *CIRCADIAN CLOCK ASSOCIATED 1* genes encode dawn-expressed, *Myb*-related repressor proteins that delay the expression of other clock genes until late in the day. Double mutant plants *(lhy cca1)* have low-amplitude, short-period rhythms that have been used in multiple studies of the plant circadian clock.

**Results**: We used *in vivo* imaging of several luciferase *(LUC)* reporter genes to test how the rhythmic gene expression of wild-type and *lhy cca1* mutant plants responded to light:dark cycles. Red, blue and red+blue light were similarly able to entrain these gene expression rhythms. The timing of expression rhythms in double mutant plants showed little or no response to the duration of light under 24h light:dark cycles (dusk sensitivity), in contrast to the wild type. As the period of the mutant clock is about 18h, we tested light:dark cycles of different duration (T cycles), simulating altered rotation of planet Earth. *lhy cca1* double mutants regained as much dusk sensitivity in 20h T cycles as the wild type in 24h cycles, though the phase of the rhythm in the mutants was much earlier than wild type. The severe, triple *lhy cca1 gi* mutants also regained dusk sensitivity in 20h cycles. The double mutant showed some dusk sensitivity under 28h cycles. *lhy cca1* double mutants under 28h cycles with short photoperiods, however, had the same apparent phase as wild-type plants.

**Conclusion:** Simulating altered planetary rotation with light:dark cycles can reveal normal circadian performance in clock mutants that have been described as arrhythmic under standard conditions. The features rescued here comprise a dynamic behaviour (apparent phase under 28h cycles) and a dynamic property (dusk sensitivity under 20h cycles). These conditional clock phenotypes indicate that parts of the clock mechanism continue to function independently of *LHY* and *CCA1*, despite the major role of these genes in wild-type plants under standard conditions.

**Accessibility:** Most results here will be published only in this format, citable by the DOI. Data and analysis are publicly accessible on the BioDare resource (www.biodare.ed.ac.uk), as detailed in the links below. Transgenic lines are linked to Stock Centre IDs below (Table 7).

## INTRODUCTION

The circadian clock allows living systems to anticipate and adapt to the day/night cycles in their environment, which are driven by the rotation of planet Earth (Millar 2016). The clock gene circuits that create and transmit biological timing are thus fundamental features of cellular physiology, in eukaryotic organisms and some prokaryotes. The clock mechanism in all organisms includes interlocked, transcriptional–translational feedback loops. The clock’s rhythmic behaviour is thought to emerge from dynamic regulation within this gene circuit, which has been well characterised in Arabidopsis (Flis et al. 2015). Figure 1 shows the normalised expression patterns of Arabidopsis clock genes under a light:dark cycle.

**Figure 1.**
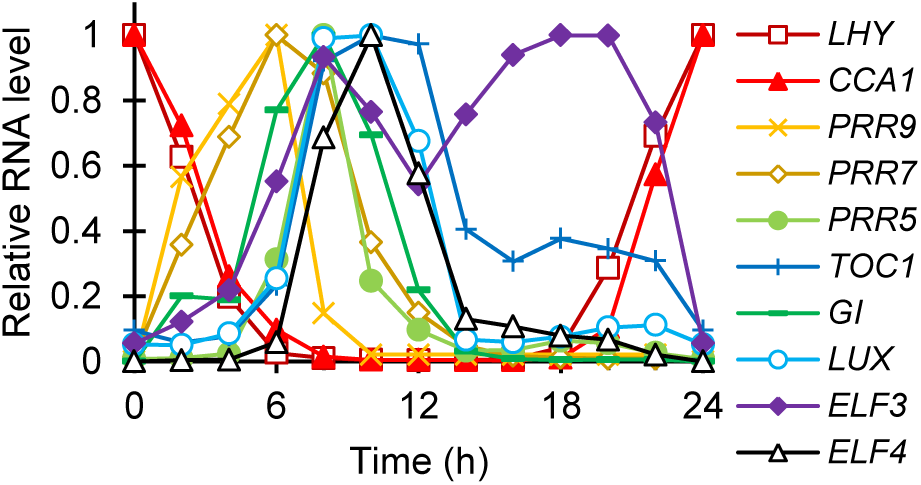
Clock gene expression in Arabidopsis. Peak-normalised RNA profiles of clock genes listed at right, in plants of the Col-0 accession under a 12h light: 12h dark cycle (LD 12:12). Time is in Zeitgeber Time (ZT, h), relative to lights-on (ZT0). In *lhy cca1* plants without the LHY and CCA1 repressors, the peak expression of all the RNAs shown here advances to ZT2-6 (data not shown). Figure adapted from (Flis et al. 2015), licensed as CC-BY.

The negative feedback loops in this model plant species incorporate two closely-related MYB transcription factors *CIRCADIAN CLOCK ASSOCIATED1 (CCA1)* and *LONG ELONGATED HYPOCOTYL (LHY)* that inhibit the expression of evening-expressed genes, such as a pseudo-response regulator (PRR) *TIMING OF CAB EXPRESSION 1 (TOC1* ≡ *PRR1)*. The expression of *CCA1* and *LHY* is tightly regulated by other clock components, including sequential inhibition by PRR9, PRR7 and PRR5. *TOC1* and other *PRR* genes are repressed by an Evening Complex, among several additional clock genes (Hsu and Harmer 2014). RVE8 (=LCL5) protein accumulates, several hours after its peak RNA abundance at dawn due to an enigmatic delay, and activates evening-expressed genes (Hsu et al. 2013), interacting with LNK proteins (Perez-Garcia et al. 2015). GI, a large plant-specific protein, is rhythmically expressed under LHY/CCA1 control but functions at a post-translational level through, for example, stabilization of the TOC1-degradation factor ZTL (Kim et al. 2007).

### The *lhy cca1* double mutant in Arabidopsis

Double mutant plants that lack both *LHY* and *CCA1* fail to repress the evening-expressed clock genes (Locke et al. 2005; Mizoguchi et al. 2002; Lu et al. 2009). The RNA levels of genes such as *TOC1* and *CCR2* (≡ *AtGRP7)* therefore rise at the start of the day, overlapping with day-phase markers such as *CAB2* (≡ *LHCB1.1)*. The amplitude of circadian rhythms in *lhy cca1* mutants damps rapidly in constant light, such that the mutants have been described as arrhythmic (Mizoguchi et al. 2002; Zeilinger et al. 2006). However, ongoing oscillations with a period of approximately 18h are reproducibly observed in double-mutant plants (Mizoguchi et al. 2002; Locke et al. 2006; Salome et al. 2010).

We earlier termed the sub-circuit that drives these, short-period rhythms the ‘Evening Loop’ and outlined its minimal properties (Locke et al. 2005). Its mechanism is only partly resolved. The rhythms in *lhy cca1* plants show that the mechanism is entrainable to 12L:12D cycles and cannot uniquely require LHY and CCA1 in order to oscillate under constant light. The long-period phenotype of *prr7 prr9* double mutants is completely suppressed by amiRNA-mediated repression of *LHY* and *CCA1* (Salome et al. 2010), indicating that PRR7 and PRR9 have no clock-relevant targets that are independent of *LHY* and *CCA1*. The rhythms of *lhy cca1* mutants are therefore unlikely to require PRR9 and PRR7. In contrast, the rhythm of *lhy cca1* plants under constant light is abolished in the triple mutants *lhy cca1 elf3* (Dixon et al. 2011) and *lhy cca1 toc1* (Ding et al.), and damps almost immediately in *lhy cca1 gi* (Locke et al. 2006). Circuit proposals including these relevant evening genes have been made in formal models (Locke et al. 2005; Pokhilko et al. 2010; Pokhilko et al. 2012; Pokhilko et al. 2013). The most recent models that show oscillation in *lhy cca1* mutants (Pokhilko et al. 2012; Pokhilko et al. 2013) depend in part upon the light-dependent dynamics of ELF3 protein degradation, mediated by COP1. *cop1* and *det1* mutants have short periods in constant light, similar to *lhy cca1* (Millar et al. 1995). The COP1 mechanism invoked is partly hypothetical (Pokhilko et al. 2011), though it has also been adopted by other researchers in this field (Shi et al. 2015). The behaviour of simulated *lhy cca1* mutants is sensitive to parameter values that are poorly constrained, however. Computational optimisation of models that include these potentially-oscillating circuits has therefore tended to lose the rhythmicity in simulated *lhy cca1* double mutants, even though the Evening Loop circuit was retained (Zeilinger et al. 2006; Fogelmark and Troein 2014).

For the clock to be useful, the endogenous period must be synchronised (entrained) to match the natural, 24-hour environmental cycle (Johnson et al. 2003). The strongest entrainment signals are temperature and light. The phase of the entrained rhythm in Arabidopsis is sensitive to multiple signals (Millar and Kay 1996; Edwards et al. 2010). Rather than tracking dawn or dusk, the clock’s phase moves earlier in shorter photoperiods (intermediate dusk sensitivity (Edwards et al. 2010)), or in entraining “T-cycles” with a period less than 24h (Somers et al.), taking several days to re-entrain (Dixon et al. 2014).

### *lhy cca1* plants as a tool to analyse starch metabolism

The remaining, partially disrupted clock circuit in *lhy cca1* double mutants allows entrainment but with strikingly altered phases. Transcripts for all the canonical clock components are expressed soon after dawn in the *lhy cca1* double mutant plants. Transcripts that peak at dawn in wild type are expressed several hours before dawn (Graf et al. 2010). The early phase of entrainment in the double mutants was used to manipulate the timing of starch degradation, which followed the predicted, early phase, and was rescued as predicted when mutant plants were tested under T=20h cycles (Graf et al. 2010). These results gave strong evidence that the starch degradation rate was set in part using the time of subjective dawn predicted by the circadian clock (Graf et al. 2010). Models based on this insight have successfully predicted starch behaviours in altered LD cycles and mutant backgrounds (Scialdone et al. 2013; Seaton et al. 2014; Pokhilko and Ebenhoh 2015).

A possible counter-argument is that starch degradation had reached some maximum limit in the double mutant, in other words the rate was constrained by other factors and was no longer responding to clock control. As noted (Seaton et al. 2014), such effects would alter the interpretation of the mutant phenotype and the rescue experiment in (Graf et al. 2010). We note that a similar effect might result indirectly, although clock control of degradation was retained, if the mutant clock was confined to an abnormal sector of phase space. In other words, the clock components oscillate in the mutant in ranges that differ from their normal values, fixing the normally rhythmic and environmentally-responsive controls on downstream processes such as the timing of starch degradation. By either direct or indirect means, under this hypothesis, the maximal starch degradation rate depleted starch to coincide quite fortuitously with subjective dawn in the mutants. The T=20h conditions shortened the night but were otherwise irrelevant. Here, we address the indirect mechanism, testing the responsiveness of the remaining clock gene circuit in the *lhy cca1* double mutant.

### Photoreceptor input to the plant clock

At least four families of photoreceptors have been identified as transducing light signals to reset the clock, the blue light sensing cryptochromes *(CRY1* and *CRY2)*, the red/far-red light (R/FR) sensing phytochromes *(PHYA, PHYB, PHYC, PHYD*, PHYE),(Devlin and Kay 2000; Somers et al. 1998a; Yanovsky et al. 2000; Edwards et al. 2015), the UV-B photoreceptor UVR8 (Feher et al. 2011) and a family of three F-box proteins, including *ZEITLUPE* (ZTL)(Baudry et al. 2010), which affect both red and blue light inputs. These eleven photoreceptors transduce light signals to regulate clock genes and proteins (Fankhauser and Staiger 2002), with both specialised and overlapping roles. It is unclear which photoreceptors mediate the entrainment of rhythms in *lhy cca1* mutant plants.

Canonical outputs from the plant clock gene circuit include highly expressed RNAs that were initially identified for their strong regulation, for example by light or cold stimuli. The promoters of *LIGHT HARVESTING COMPLEX (LHC ≡ CHLOROPHYLL A/B BINDING, CAB)* and *COLD AND CIRCADIAN REGULATED (CCR ≡ GLYCINE-RICH PROTEIN, AtGRP)* genes have been fused to firefly luciferase (*LUC*) reporter genes, to reveal the regulation of these two classes of clock-controlled genes. *In vivo* imaging of bioluminescence produced by transgenic plants containing the *LUC* reporter fusions reveals their biological rhythms (Southern and Millar 2005; van Leeuwen et al. 2000). Our earlier work used LUC reporters to measure the dusk sensitivity of entrainment in wild-type plants under conventional, 24h T-cycles with various periods (Millar and Kay 1996; Edwards et al. 2010). Here we apply these methods to the *lhy cca1* double mutant, under both normal and altered T-cycles, with varying photoperiods and light quality.

## RESULTS

### The clock in *lhy cca1* entrains to red and to blue light

Transgenic seedlings bearing the *CCR2:LUC* reporter gene were grown under white light:dark cycles (LD) and transferred to constant light, under red (R), blue (B) or R+B (R+B; equivalent to physiological ‘white’ light) LED sources. Luminescence of individual seedlings was imaged using an ultra-low-light camera. Figure 2 shows that all three light sources maintained entrainment of the rhythms at very similar phases in wild-type seedlings of the Ws accession. The expected, early phase and short period were observed in the *lhy cca1* double mutants. The phase of the mutants under LD was not significantly different in the three light qualities, though there was a tendency to earlier phase under B.

**Figure 2.**
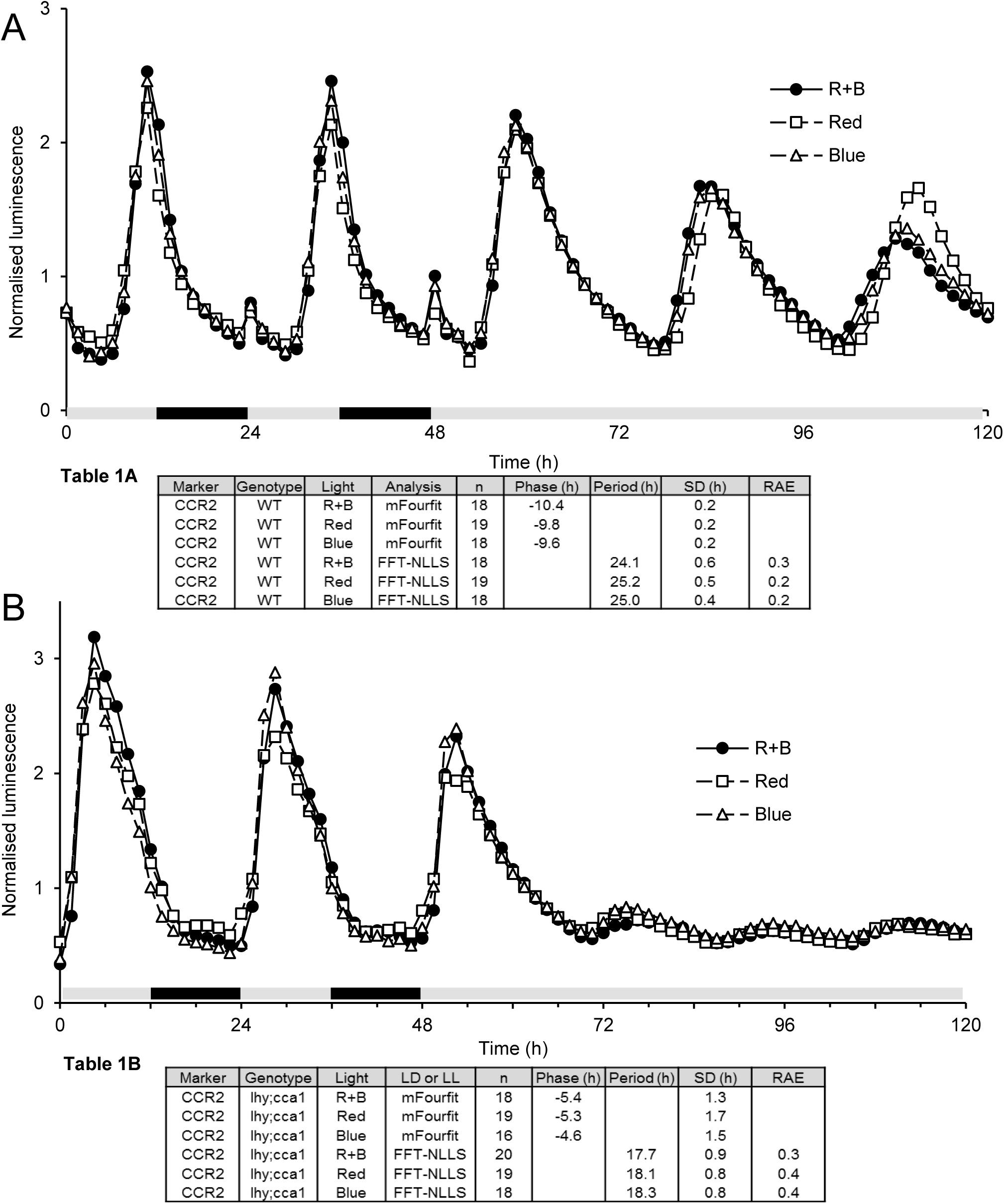
Red or blue light is sufficient to maintain entrainment of the *lhy cca1* double mutant clocks. After germination, seedlings were grown for 6 days in 12L:12D cycles of white light then transferred to the same conditions of either red, blue or red and blue light for 4 days (last 2 shown), followed by 3 days of constant light. Circadian expression of *CCR2:LUC* was tested by *in vivo* imaging in **A** wild-type plants (WT, Wassilewskija) and **B** *lhy cca1* double mutants. Grey box=light, black=dark. The data shown are genotype means, from one of 2–3 independent experiments with very similar results. **Table 1 A, B**. Phase and period analysis of data in Figure 2. The phases under LD were analysed using mFourfit; the periods of data under LL were analysed using FFT-NLLS. Periods between 15h and 35h were selected for use in **A** and **B** (n, sample number).

The apparent phase under LD cycles was estimated objectively using the mFourfit algorithm (Edwards et al. 2010), which is designed for stably entrained, non-sinusoidal waveforms (see Discussion). Period under subsequent LL were tested using FFT-NLLS (Plautz et al. 1997). Numerical results are presented in Tables 1 and 2. Table 1 shows that the phase of Ws under R+B LD was marginally later (more negative, by convention; t-test p<0.05). A phase delay in Ws during the first day of constant R delayed the peak phase by 1-1.5h relative to the B condition, though periods were not significantly different in these conditions. The mutant periods did not differ significantly. The results show that both R and B effectively entrained the double mutant, suggesting that its remaining clock circuit retains multiple light inputs.

A matching experiment was conducted under FR light (Table 2). Interpretation was hampered in some cases by low signal levels and/or amplitude from the reporter genes. Note that the behaviour of these seedlings grown without sucrose under white light and transferred to far-red light differs from the results of (Wenden et al. 2011), which used seedlings germinated under far-red light with exogenous sucrose. Nonetheless, rhythmic *GI* expression arrested close to the peak level in constant FR whereas *CCA1* expression collapsed to the trough level, as in (Wenden et al. 2011).

**Table 2.**
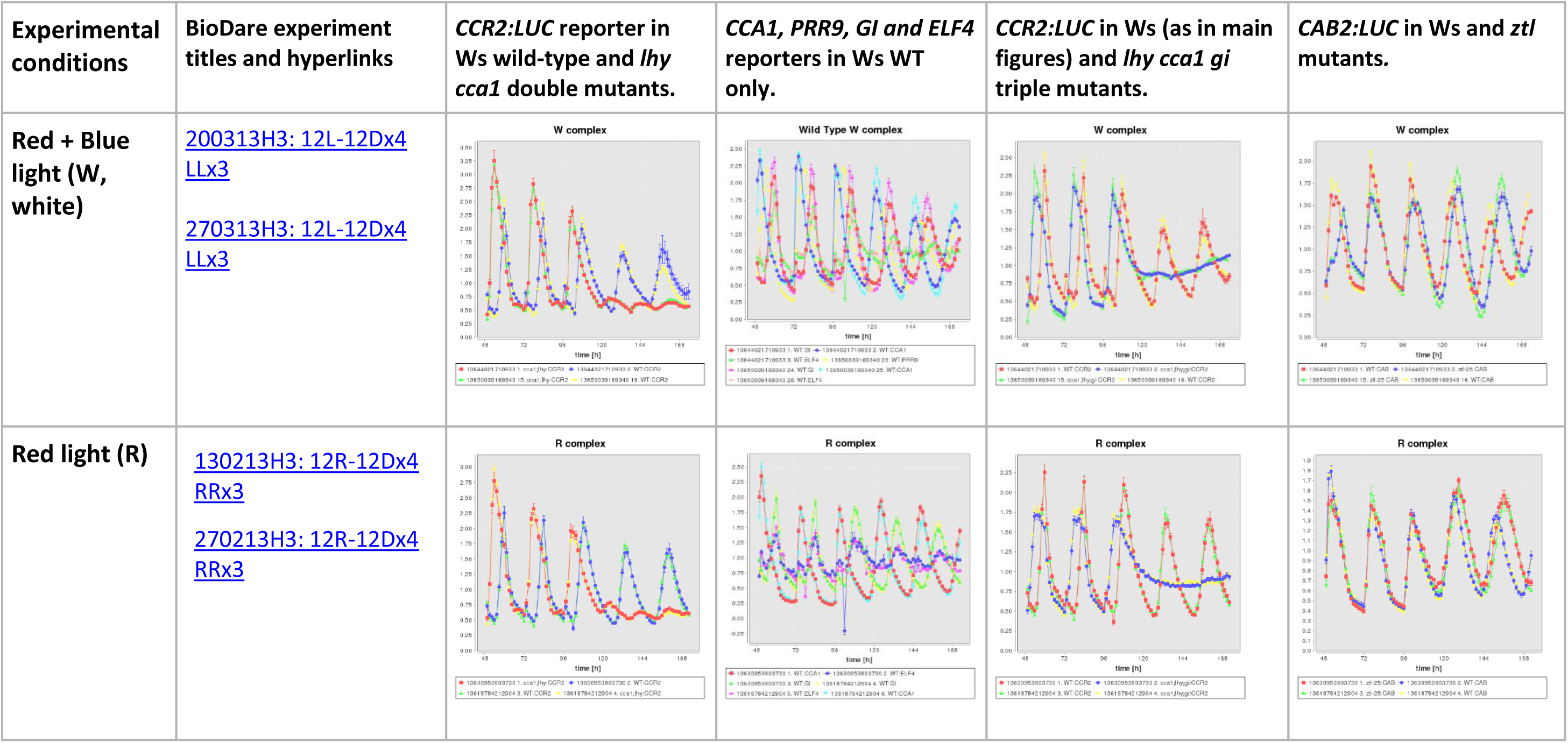
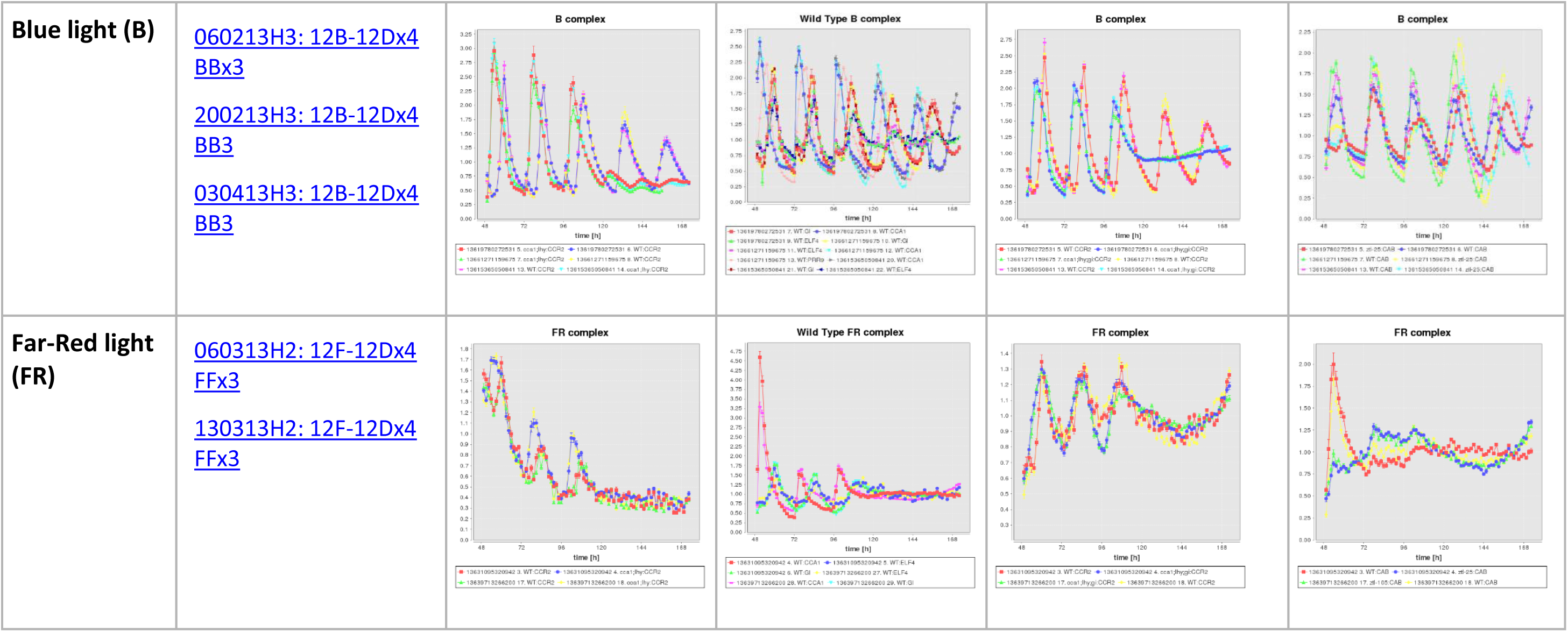
Aggregated *LUC* reporter data from all replicates of the studies summarised in Figure 2. Mean +/-SEM (n=14 to 30 plants) of the bioluminescence signal is plotted, normalised to mean of each timeseries, for the reporters and genotypes indicated in the column headers under the conditions indicated in the first column and above each plot (‘complex’ signifies a changing light condition). The 12L:12D treatments were applied at 0–96h, followed by constant light from 96h-end. Time 96h here is plotted as 48h in the main figures. The *TOC1* reporter was also tested in WT and *lhy cca1* (not shown). BioDare experiment titles and hyperlinks to the experimental records are given in the second column. *CCR2:LUC* data in Ws and *lhy cca1* are plotted without detrending, as in the main figures. The remaining plots use linear or cubic detrending. Cubic detrending can introduce an artefactual rising trend after rhythmic amplitude damps out, for example in the triple mutants. Plots were produced using the BioDare ‘Aggregate Data’ function.

### *lhy cca1* is insensitive to photoperiod under T=24h

The Arabidopsis clock has been characterised by limited dusk sensitivity. Comparing different photoperiods revealed that the time of dawn is more important than dusk in setting the phase of entrainment, though later dusk times (longer photoperiods) can delay phase by 2–4h (Millar and Kay 1996; Edwards et al. 2010). Entrainment depends upon the molecular responses of particular clock components, which rhythmically change both expression level and responsiveness. The potential molecular mechanisms for light input therefore vary over the day/night cycle. In particular, dusk sensitivity depends upon the balance of light inputs to the clock that operate in the morning and in the evening. Under standard light-dark cycles, transcripts for all the canonical clock components are expressed soon after dawn in *lhy cca1* double mutant plants. We therefore tested whether the clock’s behaviour was locked to this pattern in the mutant, altering the mutant’s dusk sensitivity and providing evidence for a restricted responsiveness to entraining cues.

We measured the phase of entrainment in response to altered entrainment regimes, using the R+B LEDs and the *CCR2:LUC* reporter. Other reporters were tested (Table 2) but *TOC1* in particular showed a more complicated rhythmic waveform that hampered phase estimation. Plants were grown and was imaged under LD cycles comprising 25%, 50% and 75% light (6L:18D, 12L:12D and 18L:6D)(Edwards et al. 2010). Expression data where the estimated period was expected for stable entrainment (24h+/-1S.D.) were selected for analysis, to avoid apparent non-stationarity due to biological noise or the analytical algorithm. Ws plants delayed the estimated phase of expression by 2.6h in 18L:6D compared with 6L:18D (12.4–12.5h after dawn compared with 9.8h after dawn; t-test p<0.03; Figure 3, Table 3), consistent with previous results (Millar and Kay 1996; Edwards et al. 2010). *lhy cca1* double mutant plants expressed the *CCR2* reporter much earlier in the day, 5.5h after dawn in short photoperiods and 5.8 to 6.2h in long photoperiods. The small, 0.3–0.7h change in apparent phase was not statistically significant. The entrainment of the double mutants under 24h cycles is therefore even less dusk-sensitive than wild-type Arabidopsis. Circadian phase in the mutants is therefore expected to track the time of dawn.

**Figure 3.**
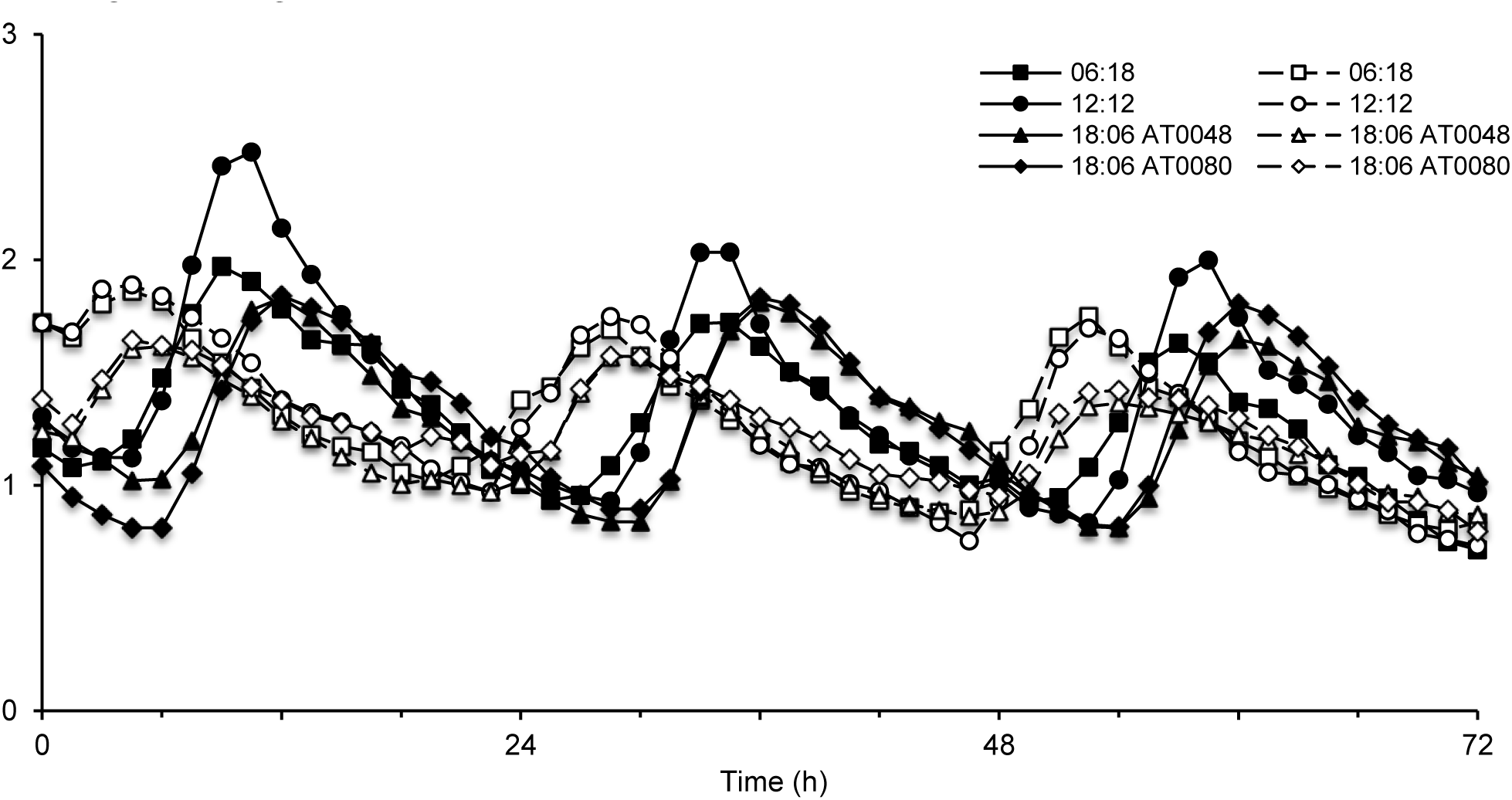
Rhythmic expression of *CCR2* in various photoperiods under T=24h cycles. A. Seedlings (n) were grown for 4, 24 hour, days (last 3 shown) under R+B light under various photoperiods; 6h light/18h dark (squares), 12h light/12h dark (circles) and 18h light/6h dark (triangles). Lights-on at 0, 24h, 48h in this timescale. This assay was carried out on both ‘WT’ (Wassilewskija, filled squares, circles, triangles and diamonds) and *cca1 lhy* double mutants (open squares, circles, triangles and diamonds). The data is representative of those analysed in Table 3. Some data for Ws were previously reported in (Edwards et al. 2010); the mutant data have not previously been published.

**Table 3.** Phase analysis of the data in **A** and **B** (1 d.p.). The phase of the final 3 days was analysed using mFourfit. Plants with estimated periods between 15 h and 35 h were tested (n samples) and those with periods of 24 h ± SD were included in the phase analysis (N samples).

| Marker | Genotype | T24 | Experiment | Analysis | n | N | Phase (h) | SD (h) |
| --- | --- | --- | --- | --- | --- | --- | --- | --- |
| CCR2 | WT | 06:18 | AT0078 | mFourfit | 21 | 19 | -9.8 | 0.7 |
| CCR2 | WT | 12:12 | AT0043 | mFourfit | 21 | 11 | -9.8 | 0.2 |
| CCR2 | WT | 18:06 | AT0048 | mFourfit | 17 | 15 | -12.4 | 0.8 |
| CCR2 | WT | 18:06 | AT0080 | mFourfit | 19 | 14 | -12.5 | 1.0 |
| CCR2 | *lhy cca1* | 06:18 | AT0078 | mFourfit | 15 | 10 | -5.5 | 2.3 |
| CCR2 | *lhy cca1* | 12:12 | AT0043 | mFourfit | 15 | 9 | -4.4 | 1.3 |
| CCR2 | *lhy cca1* | 18:06 | AT0048 | mFourfit | 6 | 6 | -5.8 | 0.7 |
| CCR2 | *lhy cca1* | 18:06 | AT0080 | mFourfit | 20 | 20 | -6.2 | 2.2 |

### *lhy cca1* adjusts phase under T=20h

In order to entrain to T=24h cycles, the short-period clock of *lhy cca1* plants must be phase-delayed by 6h (one third of its ~18h period) within every cycle. The stable phase of entrainment in the mutant is therefore very early, such that a large interval of the delaying region of the phase response curve in the early subjective night (and less of the phase-advancing region in the late night) coincides with the light interval (Johnson et al. 2003). We reasoned that this imposed, very early phase of entrainment might mask any subtler response to photoperiod that might remain possible in the double mutant. We therefore repeated the test of dusk sensitivity under T=20h cycles, much closer to the mutant’s free-running period, with LD cycles comprising 25%, 50% and 75% light (5L:15D, 10L:10D and 15L:5D; Figures 4 and 5).

**Figure 4.**
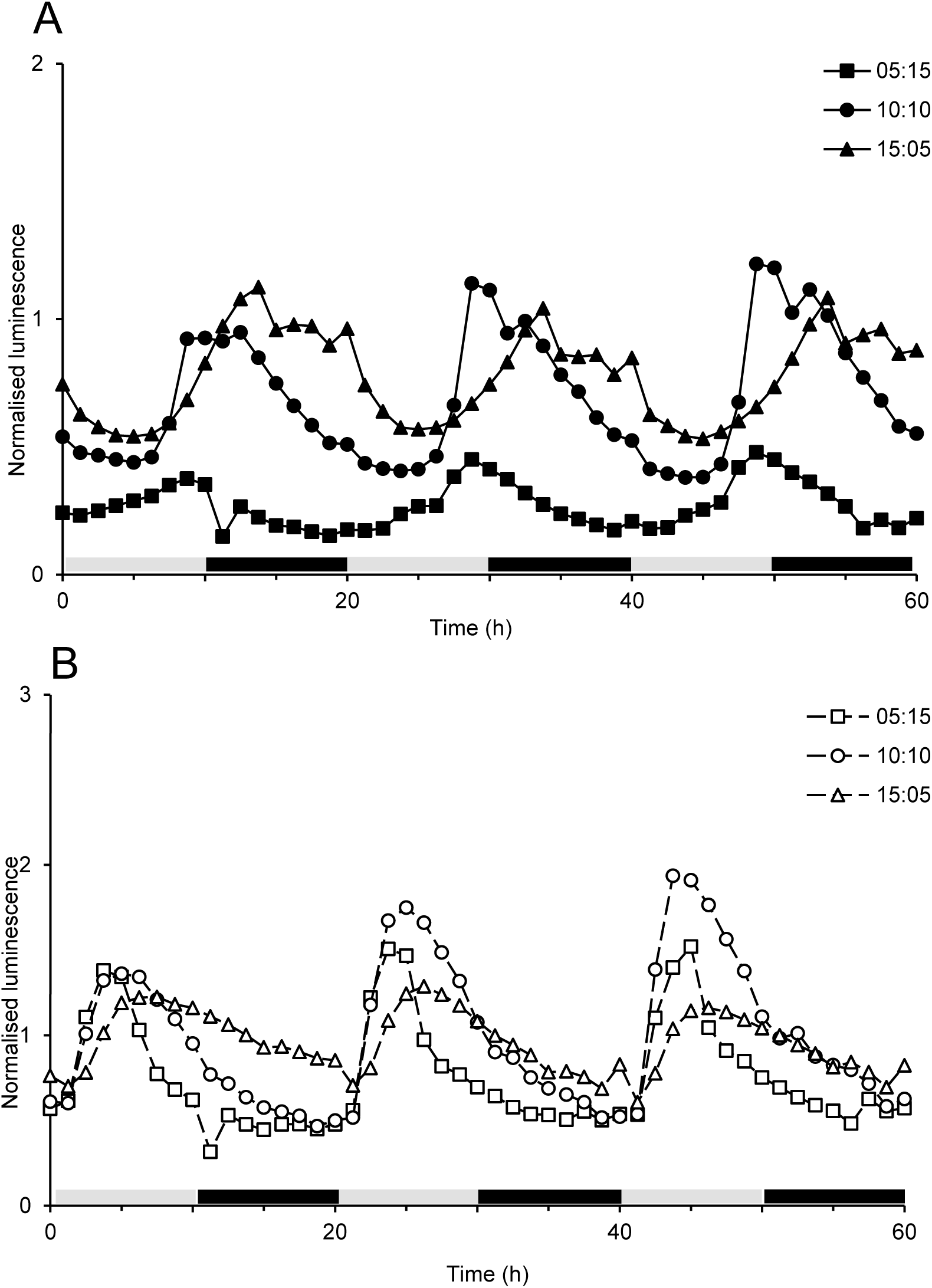
Rhythmic expression of *CCR2* exposed to various photoperiods under T=20h cycles; focused analysis. Seedlings were grown and imaged under T=20h cycles (last 3 shown) of R+B light with various photoperiods; 5h light/15h dark (squares), 10h light/10h dark (circles) and 15h light/5h dark (triangles). Data representative of those analysed in Table 4 are shown for WT, **A** (Wassilewskija, filled squares, circles and triangles) and *lhy cca1* double mutants, **B** (open squares, circles and triangles).

**Figure 5.**
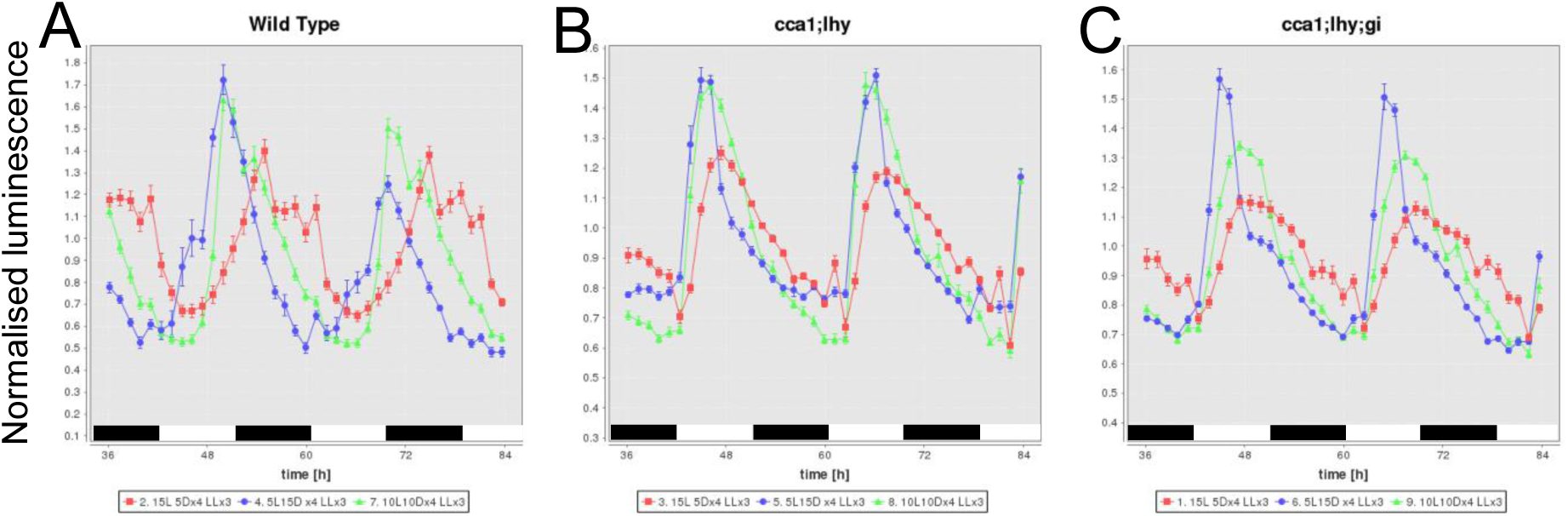
Rhythmic expression of *CCR2* exposed to various photoperiods under T=20h cycles; global analysis. The data for seedlings were grown and imaged under T=20h cycles of R+B light in figure 4 are shown for 5h light/15h dark (blue circles), 10h light/10h dark (green triangles) and 15h light/5h dark (red squares). Luminescence data were detrended in BioDare using the baseline+amplitude detrending kernel developed for mFourfit (Edwards et al. 2010) and normalized to the mean of each timeseries. Mean data +/-SEM are plotted for each reporter, genotype and condition; in the case of *lhy cca1*, these are all the data analysed in Table 5. **A**, wild-type plants (Wassilewskija); **B**, *lhy cca1* double mutants; **C**, *lhy cca1 gi* triple mutants.

**Figure 6.**
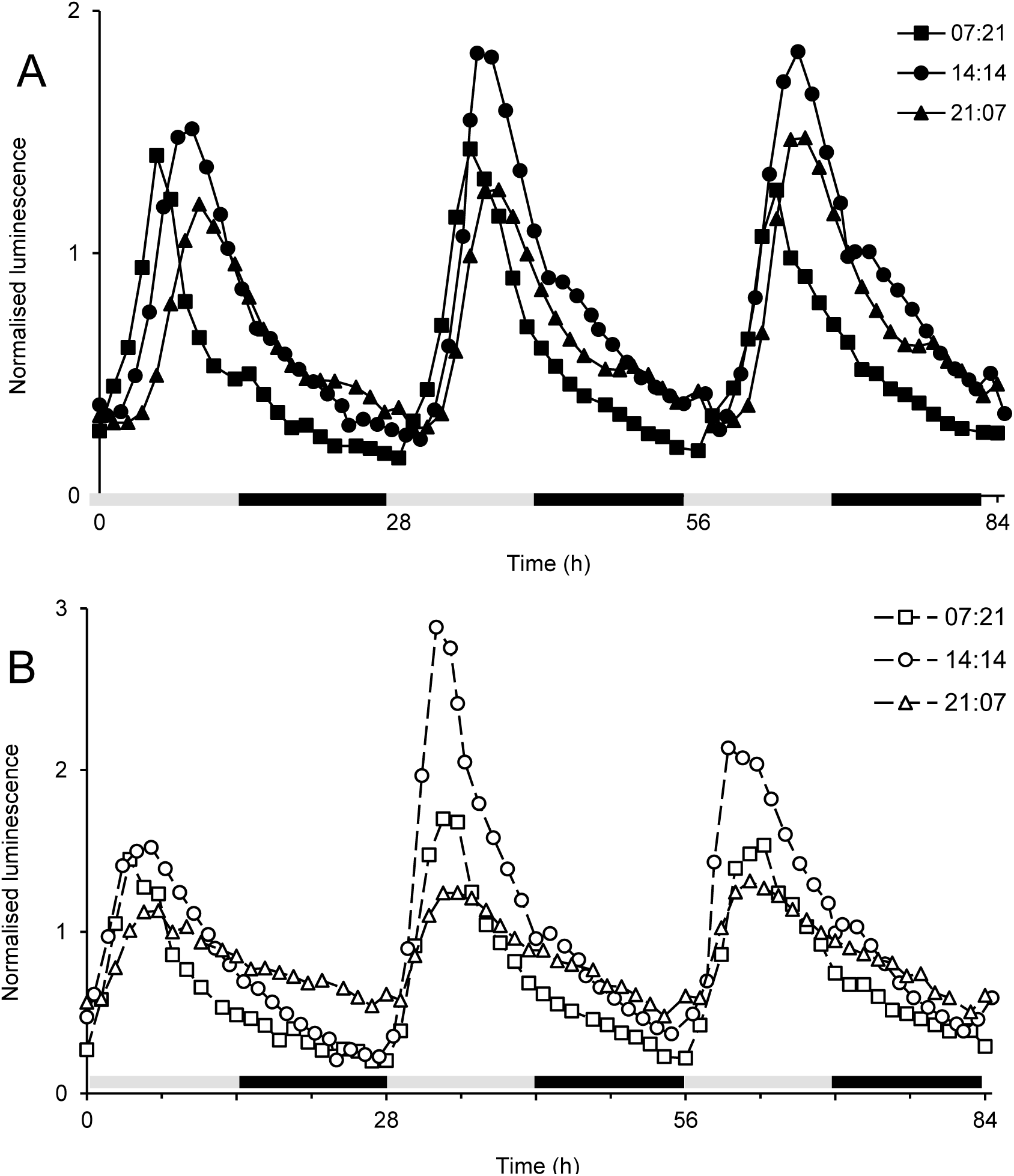
Rhythmic expression of *CCR2* in various photoperiod under T=28h cycles. Seedlings (n) were grown for 4, 28 hour, days (last 3 shown) under R+B light with various photoperiods; 7h light/21h dark (squares), 14h light/14h dark (circles) and 21h light/7h dark (triangles). This assay was carried out on both ‘WT’, **A** Wassilewskija, filled squares, circles and triangles) and *lhy cca1*, **B** double mutants (open squares, circles and triangles). The data is representative of those analysed in Table 6.

Wild-type plants must phase-advance by 4h or more within each cycle, in order to entrain to T=20h. The stable phase of entrainment is therefore expected to be late in the WT, so that a sufficiently large phase-advancing region around subjective dawn coincides with the light interval (Johnson et al. 2003). This effect was apparently small: the estimated phase under 10L:10D was 10.2h after dawn, similar to the estimated phase under 12L:12D. The estimated phase of WT plants was 1.6h later in 10h compared to 5h photoperiods (Table 4); altered waveform complicated phase estimation under 15h photoperiods but visual estimates suggested a 9h phase difference in 15h compared to 5h photoperiods.

Expression patterns in the *lhy cca1* mutants were similar under 5L:15D and 10L:10D. The peak times remain much earlier in the double mutant than in the wild type, such that the phase difference from wild type was greatest in 15L:5D (>7h, data not shown; the complicated waveform in WT hampers the comparison). The peak phase was delayed in *lhy cca1* by 2.3–2.7h in 15L:5D relative to the shorter photoperiods (Tables 4, 5), in contrast to their behaviour under T=24h cycles. This result was confirmed by re-analysis of the data using different phase estimates, without pre-selecting fitted periods close to 24h, and with a radically different analytical algorithm (Enright periodogram; Table 5). Thus, the remaining clock gene circuit in the double mutants can respond to photoperiod under T=20h, with the same dusk sensitivity as a WT plant under T=24h.

Unexpectedly, the apparent phase of the *lhy cca1 gi* triple mutant responded even more than in *lhy cca1* (Figure 5C). Under constant conditions, rhythms in the triple mutant usually damp after only one cycle (Locke et al. 2006), indicating that its clock defect is even more severe than in *lhy cca1*. However, *lhy cca1* de-represses evening gene expression including expression of *GI*. Removing GI function might partially restore some aspects of regulation in the triple mutant. Millar et al. 2015

**Table 4.** Phase analysis of the data in **A** and **B** (1 d.p.). The phase of the final 3 days was analysed using mFourfit. Periods between 15 h and 35 h were selected for use in **A** and **B** (n) and fits within the range of the expected period 24 h ± SD were used for the phase analysis (N).

| Marker | Genotype | T20 | Experiment | Analysis | n | N | Phase (h) | SD (h) |
| --- | --- | --- | --- | --- | --- | --- | --- | --- |
| *CCR2* | WT | 05:15 | 060313H1 | mFourfit | 20 | 12 | -8.6 | 0.6 |
| *CCR2* | WT | 10:10 | 270213H1 | mFourfit | 20 | 12 | -10.2 | 1.5 |
| *CCR2* | WT | 15:05 | 290313H2 | mFourfit | 17 | 16 | 10.3 | 4.0 |
| *CCR2* | *lhy cca1* | 05:15 | 060313H1 | mFourfit | 20 | 15 | -4.3 | 0.2 |
| *CCR2* | *lhy cca1* | 10:10 | 270213H1 | mFourfit | 17 | 12 | -4.7 | 0.3 |
| *CCR2* | *lhy cca1* | 15:05 | 290313H2 | mFourfit | 21 | 19 | -7.0 | 1.5 |

**Table 5.** Phase analysis of the data in **A** and **B** (1 d.p.). The phase of the final 3 cycles of data was analysed after linear detrending, using mFourfit (BioDare analysis jobs 7501 and 7503) and the Enright periodogram (jobs 7502 and 7504). All periods between 15 h and 25 h were used for the phase analysis (N). Group statistics are reported with (Weighted) and without (Arithmetic) weighting by BioDare’s Goodness-of-Fit metric.

| Genotype | Marker | Conditions | Algorithm | N | Weighted phase |  |  | t-test p | Arithmetic phase |  |  | t-test p |
| --- | --- | --- | --- | --- | --- | --- | --- | --- | --- | --- | --- | --- |
|  |  |  |  |  | Mean | SE | SD |  | Mean | SE | SD |  |
| *lhy cca1* | *CCR2* | 5L:15D | mFourfit | 20 | 1.3 | 0.08 | 0.38 | <0.0001 | 1.3 | 0.11 | 0.49 | <0.01 |
| *lhy cca1* | *CCR2* | 15L:5D | mFourfit | 20 | 4.0 | 0.48 | 2.17 |  | 4.9 | 0.71 | 3.17 |  |
| *lhy cca1* | *CCR2* | 5L:15D | ER Periodogram | 20 | 0.8 | 0.10 | 0.46 | <0.0001 | 0.7 | 0.12 | 0.55 | <0.01 |
| *lhy cca1* | *CCR2* | 15L:5D | ER Periodogram | 20 | 3.6 | 0.51 | 2.28 |  | 2.8 | 0.73 | 3.25 |  |
| *lhy cca1* | *TOC1* | 5L:15D | mFourfit | 19 | 1.5 | 0.08 | 0.34 | <0.0001 | 1.5 | 0.08 | 0.37 | <0.0001 |
| *lhy cca1* | *TOC1* | 15L:5D | mFourfit | 20 | 4.0 | 0.24 | 1.06 |  | 4.3 | 0.46 | 2.06 |  |
| *lhy cca1* | *TOC1* | 5L:15D | ER Periodogram | 19 | 1.0 | 0.14 | 0.60 | <0.0001 | 1.1 | 0.16 | 0.68 | <0.0001 |
| *lhy cca1* | *TOC1* | 15L:5D | ER Periodogram | 20 | 4.4 | 0.29 | 1.31 |  | 4.6 | 0.38 | 1.68 |  |

### Both wild-type and *lhy cca1* plants increase response to photoperiod under T=28h

The severity and pleiotropy of phenotypes in the *lhy cca1* mutant might alter circadian entrainment by multiple, indirect mechanisms. The limited dusk sensitivity of the double mutants observed under T=24h (Figure 2) might therefore be a mutant-specific effect, rather than a general property of the plant circadian circuit when entrained to a T-cycle longer than its free-running period. To control for such effects, we tested the dusk sensitivity of wild-type plants under T=28h, which must also delay phase by ~4h every cycle in order to entrain. The phases observed were advanced 2–3h (Table 6) relative to the phases under equivalent T=24h cycles (Table 3). This is consistent with the expected requirement for more of the early subjective night to coincide with the light interval. WT plants showed a 4h delay in 21L:7D cycles relative to 7L:21D, greater than in T=24h cycles. *lhy cca1* mutant plants had early phases, as expected. The mutants also showed a larger phase delay than in T=24h; the change between 7 and 21h photoperiods was 1.7h, which was statistically significant (t-test p<0.001). Strikingly, the early phase of the wild-type plants nearly synchronized their *CCR2* expression with the double mutants, especially in 7L:21D (Figure 7B). Thus the timing of *CCR2* expression can be independent of *LHY* and *CCA1*. The data also suggested a 24h period component in constant light in *lhy cca1*, albeit at very low amplitude (Figure 7C), which we have not previously observed.

The long-period *ztl* mutant (tau=28h) was also tested under each condition using the *CAB2* reporter, to test whether a long-period mutant that was forced to phase-advance could lose dusk sensitivity (data not shown; available on BioDare, please see Data Accessibility).

**Table 6.** Phase analysis of the data in **A** and **B** (1 d.p.). The phase of the final 3 days was analysed using mFourfit. Periods between 15 h and 35 h were selected for use in **A** and **B** (n) and the range of the expected period 24 h ± SD was used for the phase analysis (N).

| Marker | Genotype | T28 | Experiment | Analysis | n | N | Phase (h) | SD (h) |
| --- | --- | --- | --- | --- | --- | --- | --- | --- |
| CCR2 | WT | 07:21 | 200313H2 | mFourfit | 15 | 8 | -6.2 | 0.6 |
| CCR2 | WT | 14:14 | 130313H1 | mFourfit | 18 | 14 | -8.8 | 0.4 |
| CCR2 | WT | 21:07 | 270313H1 | mFourfit | 20 | 13 | -10.2 | 1.2 |
| CCR2 | *lhy cca1* | 07:21 | 200313H2 | mFourfit | 20 | 13 | -5.1 | 0.7 |
| CCR2 | *lhy cca1* | 14:14 | 130313H1 | mFourfit | 18 | 7 | -5.5 | 0.8 |
| CCR2 | *lhy cca1* | 21:07 | 270313H1 | mFourfit | 17 | 7 | -6.8 | 0.9 |

## DISCUSSION

Circadian clock mechanisms include gene regulation by multiple, interlocking feedback loops, which can increase the flexibility of possible regulatory changes over evolutionary time and in the face of environmental variations. In light-grown Arabidopsis seedlings, multiple photoreceptors contribute light input signals, adding further complexity to the clock network. The interaction of light input and clock mechanism that normally leads to dusk-sensitive entrainment in the wild-type plants under T=24h cycles was disrupted in the *lhy cca1* double mutant, reducing or eliminating its dusk sensitivity under T=24 and T=28h cycles. We showed that 20h cycles rescued dusk sensitivity in the apparent phase of the *lhy cca1* mutant.

Circadian phase is commonly tested by measuring rhythms under constant conditions, without the masking effects of the light:dark transitions, extrapolating back to the phase at the time of the last entraining stimulus. This extrapolation proved impossible, owing to the drastic drop in amplitude and period of *lhy cca1* plants between LD and LL (Figure 2B). Apparent phase during light:dark cycles will be affected by direct and indirect light regulation of the reporter. The *CCR2* fusion was selected because it showed fewer such effects than the other markers, in particular compared to the *TOC1* reporter that was also introduced into the *lhy cca1* mutant (Locke et al. 2006).

Our results suggest that the expression of the remaining clock genes in *lhy cca1* mutants is not simply forced by overwhelming regulation from the light:dark cycle, locking the rhythms into a phase close to dawn. Under T=20h cycles in particular, the clock reporter achieved graded, photoperiod-dependent control and thus regained dusk-sensitive entrainment. There is therefore no reason to expect that downstream processes will be locked into maximal or minimal states either, nor in particular, that clock control of starch degradation would be locked at its maximal rate under 10L:10D cycles as in (Graf et al. 2010). Indeed, our data suggest that even *lhy cca1* plants might modulate the rate of starch degradation in response to changing photoperiods under T=20h cycles, given that they can alter the phase of biological rhythms in these conditions. The RNA expression levels of clock genes also remained within their normal range in *lhy cca1* under LD cycles, despite the change in their timing (Flis et al. 2015). However, the phase of *CCR2* expression relative to the 10L: 10D cycle was advanced more than in wild-type plants under 12L:12D (Figures 3, 4B, 5B), whereas the timing of starch degradation was restored close to its normal timing under 10L:10D (Graf et al. 2010). Therefore *CCR2* might not be an ideal marker for the (unknown) subjective dawn predictor that is used to control starch degradation (Scialdone et al. 2013; Seaton et al. 2014). Indeed, the *GBSS* RNA was used in (Graf et al. 2010) as a marker for subjective dawn, as it was expressed in phase with *LHY* and *CCA1* in the wild type.

Finally, our results show that aspects of circadian timing under different conditions can be surprisingly independent of *LHY* and *CCA1*. The wild-type and mutant expression profiles differed most under long photoperiods in short, T=20h cycles (Figure 7A), but were strikingly similar under short photoperiods in long, T=28h cycles (Figure 7B). These conditions require opposite phase shifts for the clocks to entrain, so it is not surprising that different clock components are involved. This approach to define discriminating conditions that reflect functionally distinct aspects of the clock mechanism was best illustrated in the ‘circadian resetting surface’ defined for *N. crassa* (Remi et al. 2010). Some of the conditions defined here will likely reveal new aspects and interactions in the clock mechanism, if a more comprehensive set of clock RNA markers is tested in future (as in Flis et al. 2015), although the assays are more laborious than the *LUC* reporters tested here.

**Figure 7.**
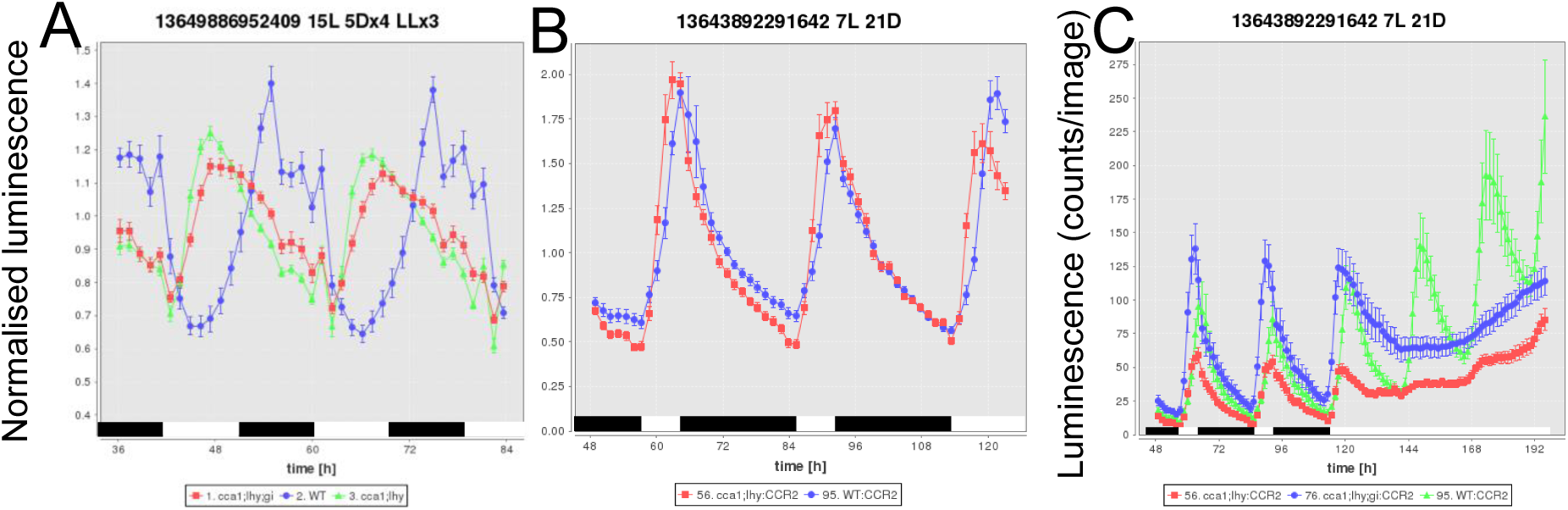
Conditional severity of the *lhy cca1* mutant phenotype. *CCR2:LUC* data are replotted, **A** from Figure 5, WT (blue circles), *lhy cca1* (green triangles) and *lhy cca1 gi* (red squares) under T=20h cycles of 15h light/5h dark, where the mutants are most different from wild type; **B**, from Figure 6, WT (blue circles), *lhy cca1* (green triangles) and *lhy cca1 gi* (red squares) under T=20h cycles of 15h light/5h dark, where the mutants are most different from wild type; **C**, as in B but showing the data in subsequent constant light: WT (green triangles), *lhy cca1* (red squares) and *lhy cca1 gi* (blue circles). Luminescence data were detrended in BioDare using the baseline+amplitude detrending kernel developed for mFourfit (Edwards et al. 2010) (A), cubic detrending (B) or no detrending (C), with (A,B) or without (C) normalisation to the mean of each timeseries. Mean data +/-SEM are plotted for each reporter, genotype and condition.

## DATA ACCESSIBILITY

The accessibility of resources used in the publication is summarised at the University of Edinburgh’s institutional repositories with the following URLs: http://www.research.ed.ac.uk/portal/en/datasets/data-sets-for-millar-et-al-biorxiv-2015(80674a78–2140–45ed-9972–3beebaf98024).html

The data sets reported here are publicly available from BioDare with the permanent data identifiers listed below, using login name ‘public’ with password ‘public’. In addition to the numerical data and analytical results, several other reporters, genotypes and replicates tested in the same studies but not published here are included in these links.

**Figure 1** shows normalised data from the following data set:

A. Flis, V. Mengin, R. Sulpice and M. Stitt (2015) TiMet RNA timeseries data from rosette plants for clock model parameterisation. Experiment TiMEt ros, BioDare accession 2841,

https://www.biodare.ed.ac.uk/robust/ShowExperiment.action?experimentId=2841

**Figure 2, Table 2:**

J.T. Carrington, S.K. Hodge and A.J. Millar (2013) Luciferase reporter timeseries data under 12L:12D cycles of red (R) light; experiment 270213H3, BioDare ID 13630953803730;

https://www.biodare.ed.ac.uk/robust/ShowExperiment.action?experimentId=13630953803730

Replicated in experiment 130213H3: 12R-12Dx4 RRx3, BioDare ID 13618784212904;

https://www.biodare.ed.ac.uk/robust/ShowExperiment.action?experimentId=13618784212904.

J.T. Carrington, S.K. Hodge and A.J. Millar (2013) Luciferase reporter timeseries data under 12L:12D cycles of blue (B) light; experiment 200213H3, BioDare ID 13619780272531,

https://www.biodare.ed.ac.uk/robust/ShowExperiment.action?experimentId=13619780272531

Replicated in experiments:

060213H3, BioDare ID 13615365050841, https://www.biodare.ed.ac.uk/robust/ShowExperiment.action?experimentId=13615365050841

030413H3, BioDare ID 13661271159675, https://www.biodare.ed.ac.uk/robust/ShowExperiment.action?experimentId=13661271159675

J.T. Carrington, S.K. Hodge and A.J. Millar (2013) Luciferase reporter timeseries data under 12L:12D cycles of red + blue (R+B) light; experiment 270313H3, BioDare ID 13650039169340, https://www.biodare.ed.ac.uk/robust/ShowExperiment.action?experimentId=13650039169340

Replicated in experiment 200313H3, BioDare ID 13644021710933, https://www.biodare.ed.ac.uk/robust/ShowExperiment.action?experimentId=13644021710933

Matching data under far-red light, Table 2 only:

J.T. Carrington, S.K. Hodge and A.J. Millar (2013) Luciferase reporter timeseries data under 12L:12D cycles of far-red (FR) light; experiment 060313H2, BioDare accession 13631095320942, https://www.biodare.ed.ac.uk/robust/ShowExperiment.action?experimentId=13631095320942

Replicated in experiment 130313H2, BioDare ID 13639713266200, https://www.biodare.ed.ac.uk/robust/ShowExperiment.action?experimentId=13639713266200

**Figure 3**

K.D. Edwards, A. Thomson and A.J.Millar (2007) Luciferase reporter timeseries data under 6L:18D cycles of red+blue light; experiment AT0078, BioDare ID 12729899933214, https://www.biodare.ed.ac.uk/robust/ShowExperiment.action?experimentId=12729899933214

K.D. Edwards, A. Thomson and A.J.Millar (2006) Luciferase reporter timeseries data under 12L:12D cycles of red+blue light; experiment AT0043, BioDare ID 12730739255199, https://www.biodare.ed.ac.uk/robust/ShowExperiment.action?experimentId=12730739255199

K.D. Edwards, A. Thomson and A.J.Millar (2006) Luciferase reporter timeseries data under 18L:6D cycles of red+blue light; experiment AT0048, BioDare ID 12730752301671, https://www.biodare.ed.ac.uk/robust/ShowExperiment.action?experimentId=12730752301671

K.D. Edwards, A. Thomson and A.J.Millar (2007) Luciferase reporter timeseries data under 18L:6D cycles of red+blue light; experiment AT0080, BioDare ID 12730527312073, https://www.biodare.ed.ac.uk/robust/ShowExperiment.action?experimentId=12730527312073

**Figures 4 and 5**

W.V. Tee, S.K. Hodge and A.J. Millar (2013) Luciferase reporter timeseries data under 5L:15D cycles of red+blue light; experiment 060313H1, BioDare ID 13631017726171, https://www.biodare.ed.ac.uk/robust/ShowExperiment.action?experimentId=13631017726171

Replicated partially in experiment 130212H1, BioDare accession 13615471409903, https://www.biodare.ed.ac.uk/robust/ShowExperiment.action?experimentId=13615471409903

W.V. Tee, S.K. Hodge and A.J. Millar (2013) Luciferase reporter timeseries data under 10L:10D cycles of red+blue light; experiment 270213H1, BioDare accession 3624982192162, https://www.biodare.ed.ac.uk/robust/ShowExperiment.action?experimentId=13624982192162

W.V. Tee, S.K. Hodge and A.J. Millar (2013) Luciferase reporter timeseries data under 15L:5D cycles of red+blue light; experiment 290313H2, BioDare accession 13649886952409, https://www.biodare.ed.ac.uk/robust/ShowExperiment.action?experimentId=13649886952409

Replicated partially (WT incomplete entrainment) in experiment 200213H1, BioDare accession 13618962781900, https://www.biodare.ed.ac.uk/robust/ShowExperiment.action?experimentId=13618962781900

**Figure 6**

W.V. Tee, S.K. Hodge and A.J. Millar (2013) Luciferase reporter timeseries data under 7L:21D cycles of red+blue light; experiment 200313H2, BioDare accession 13643892291642, https://www.biodare.ed.ac.uk/robust/ShowExperiment.action?experimentId=13643892291642

W.V. Tee, S.K. Hodge and A.J. Millar (2013) Luciferase reporter timeseries data under 14L:14D cycles of red+blue light; experiment 130313H1, BioDare accession 13637871113032, https://www.biodare.ed.ac.uk/robust/ShowExperiment.action?experimentId=13637871113032

W.V. Tee, S.K. Hodge and A.J. Millar (2013) Luciferase reporter timeseries data under 21L:7D cycles of red+blue light; experiment 270313H1, BioDare accession 13649842953637, https://www.biodare.ed.ac.uk/robust/ShowExperiment.action?experimentId=13649842953637

## EXPERIMENTAL PROCEDURES

### Plant material

The transgenic Arabidopsis lines carrying *CCR2:LUC* reporters in the Ws background have been described previously (Locke et al. 2006) and several are available from the community stock centers (Table 7).

### Growth conditions and LUC imaging

Growth conditions were similar to those described for Figure 3 (Edwards et al. 2010). The seeds were plated on sterile media in 12cm x 12cm square tissue culture plates, comprising 0.5x Murashige and Skoog salts (Melford M221; 0.215% m/v) with 1.2% (m/v) agar (Sigma A1296–500G), pH 5.8. Each plate contained 3 transgenic lines with 30 seeds per line. The plates were sealed with microporous tape and stratified at 4°C in the dark for 4–5 days. For Figure 2, seeds were germinated in a Sanyo MLR-351H Plant Growth Chamber at 22°C under white fluorescent light (75 μmol.m^-2^. s^-1^) for 6 days of 12L:12D, before transfer to the experimental lighting conditions in Percival I-30BLL cabinets illuminated by custom-made LEDs (Nipht, Edinburgh, Scotland). The total light intensity in the imaging cabinet was 25μmol.m-2.s-1 for each light quality. For figures 4–7, seeds were grown under white fluorescent light in the experimental photoperiod and T-cycle for 6 days. On the sixth day the seedlings were sprayed with 5mM D-luciferin and 0.01% Triton X-100 solution and transferred to the imaging cabinet, where experimental conditions were maintained with red (72 μmol m-2s-1) and blue (40 μmol m-2s-1) LEDs (Nipht, Edinburgh). A Percival I-30BLL cabinet and Hamamatsu C4742–98 digital camera operated by Wasabi software (Hamamatsu Photonics, Hamamatsu City, Japan) were used for data acquisition, illuminated by custom-made LEDs (Nipht, Edinburgh, Scotland). 4 dishes were imaged in each camera. Each condition for Figure 2 was replicated 2–3 times. The larger number of conditions in Figures 4–7 were tested only once each, with incomplete replicates of 5L:15D and 15L: 5D.

### Measurement of circadian rhythms

Individual period and phase estimates were produced from bioluminescence data essentially as described (Edwards et al. 2010). Total luminescence per seedling and timepoint were determined from the image stacks after subtracting background from a ‘cone of darkness’ located between the four plates, using Metamorph software. Circadian phase of each seedling during LD cycles was estimated using the mFourfit algorithm and (in Table 5) the Enright periodogram, circadian period in LL was estimated using the FFT-NLLS algorithm (Zielinski et al. 2014). All analysis methods were accessed through the BioDare online resource (Moore et al. 2014), where all the numerical data and analytical results are publicly available along with several other reporters, genotypes and replicates tested in the same studies but not published here (please see Data Accessibility section). Group statistics were variance-weighted except for Table 5, which used weighting by BioDare’s Goodness-of-Fit metric.

The Goodness of Fit (GOF) is designed to be independent of the rhythm analysis method, allowing multiple methods to be compared. GOF is defined as the ratio of two errors. The method fit error is the error between the original time series data and the curve predicted by the rhythm analysis method. The polynomial fit error is the error between the original time series data and a non-rhythmic, polynomial (cubic) curve fitted to the time series. The GOF ratio can vary from 0 (model provides a perfect fit to the data) to a large number, indicating that the model is no better than (or is worse than) a cubic fit to the data.

**Table 7.**
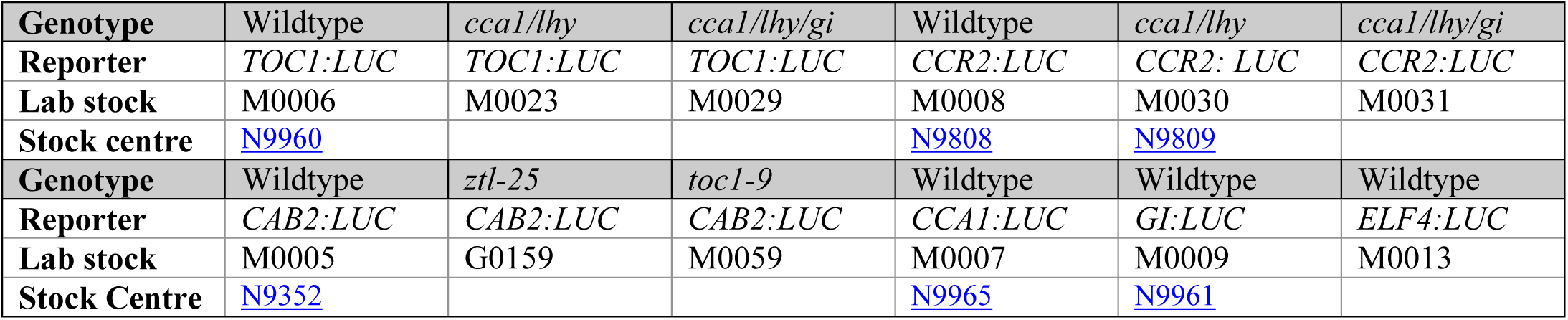
Stock centre and lab designations of transgenic lines used in the experiments under R and B, T20 and T28. Data for wild-type plants and *lhy cca1* double mutants are presented in this paper. Experiments under T24 in Figure 2 used earlier stocks of the same transgenic lines (Edwards et al. 2010).

| Genotype | Wildtype | *cca1/lhy* | *cca1/lhy/gi* | Wildtype | *cca1/lhy* | *cca1/lhy/gi* |
| --- | --- | --- | --- | --- | --- | --- |
| Reporter | *TOC1:LUC* | *TOC1:LUC* | *TOC1:LUC* | *CCR2:LUC* | *CCR2: LUC* | *CCR2:LUC* |
| Lab stock | M0006 | M0023 | M0029 | M0008 | M0030 | M0031 |
| Stock centre | N9960 |  |  | N9808 | N9809 |  |
| Genotype | Wildtype | *ztl-25* | *toc1-9* | Wildtype | Wildtype | Wildtype |
| Reporter | *CAB2:LUC* | *CAB2:LUC* | *CAB2:LUC* | *CCA1:LUC* | *GI:LUC* | *ELF4:LUC* |
| Lab stock | M0005 | G0159 | M0059 | M0007 | M0009 | M0013 |
| Stock Centre | N9352 |  |  | N9965 | N9961 |  |

## Acknowledgements

We are grateful to Kieron Edwards and Adrian Thomson for performing the experiments shown in Figure 3, to Tomasz Zielinski for the online BioDare resource and to members of the Millar laboratory for data curation in BioDare.

